# Brain predictive models of cognition fail to generalize across ethnicities: Modality-dependent bias in MRI-based prediction

**DOI:** 10.1101/2025.11.12.688133

**Authors:** Farzane Lal Khakpoor, William van der Vliet, Jeremiah Deng, Narun Pat

**Affiliations:** Department of Psychology, University of Otago, Dunedin, New Zealand; School of Computing, University of Otago, Dunedin, New Zealand

**Keywords:** neuroimaging, predictive modelling, bias, cognitive prediction, population generalizability, model fairness

## Abstract

Predictive neuroimaging models promise precision medicine but risk exacerbating health inequities if they perform unevenly across ethnic/racial groups. Using the Adolescent Brain Cognitive Development data, we benchmarked ethnic/racial bias in models predicting cognitive functioning from 91 MRI phenotypes across four training strategies. Models trained on one ethnicity performed best within that group. Models trained on participants sampled without regard to ethnicity, a common practice, performed better on White participants, likely because the ABCD sample was predominantly White. Training on equal-sized White and African American subsamples reduced disparities without accuracy loss, emerging as the upper bound for both accuracy and fairness. Structural MRI exhibited the greatest bias, whereas task-based fMRI phenotypes were more equitable. Stronger brain–cognition associations generalized more equitably, but multimodal stacking—despite enhancing prediction—did not improve fairness. Increasing representation of African American participants improved performance up to balanced sampling, with diminishing returns beyond. This first modality-wide benchmark reveals pervasive, modality-dependent ethnic bias in cognitive prediction and identifies key factors shaping equity in neuroimaging models.

## Introduction

The use of machine learning in predictive modelling enables neuroscientists to move beyond associating individual brain features to variables of interest, such as cognitive functioning, toward predicting these variables in new unseen participants or patients based on high-dimensional brain features. This approach has accelerated biomarker discovery and paved the way for the development of precision medicine [1] with the potential to support earlier risk detection and more targeted intervention [2]. For instance, predictive modelling has transformed how we study individual differences in cognitive functioning [3–6], a core psychiatric phenotype that is strongly associated with daily functioning and treatment outcomes [7]. Despite these advances, a critical translational challenge remains: ensuring that predictive models generalise and perform consistently across diverse populations.

Ethnic/racial bias in predictive models is well-documented across multiple biomedical domains [8–11]. In genomics, for instance, polygenic risk scores trained predominantly on European-ancestry cohorts often exhibit reduced accuracy in individuals from other ancestries. This illustrates how the composition of a training cohort, referred to as the discovery group in genomics, can systematically disadvantage underrepresented populations [12–14]. Such disparities in predictive performance raise concerns that these tools may inadvertently exacerbate existing health inequities [15,16].

Neuroimaging could face similar challenges. Large-scale imaging datasets are disproportionately composed of White participants from North America and Europe [17], and models trained on such data may learn associations that are less stable or less relevant in underrepresented populations. Recent work has begun to quantify this risk. For example, Li and Colleagues [18] reported that resting-state functional connectivity models trained on majority White American samples from the Adolescent Brain Cognitive Development (ABCD) study generalized poorly to African American participants, typically favouring White Americans in predicting a range of cognitive and behavioural measures. Another recent study found that brain-age algorithms based on structural data exhibit lower accuracy among African American participants [19]. These findings underscore that ethnic/racial biases are not confined to genomics but are also relevant to neuroimaging-based prediction.

However, several open questions remain. First, the scale and scope of ethnic/racial bias in predictive neuroimaging are not well characterized. It is unclear whether poor cross-ethnicity generalizability is confined to specific MRI-derived phenotypes or extends across the full spectrum of neuroimaging types. Different MRI-based representations—ranging from task-related contrasts to connectivity and structural phenotypes—may vary in their bias and susceptibility to training sample composition, such as models trained on a single ethnicity, balanced datasets including equal samples of each ethnicity, or imbalanced majority–minority combinations [18]. A systematic benchmarking across neuroimaging phenotypes is therefore needed to determine which representations are more or less sensitive to training composition and which yield more equitable predictions.

Second, the factors influencing such disparities remain poorly understood. Given that neuroimaging phenotypes differ in their predictive utility [20–22], it is important to determine whether neuroimaging phenotypes that capture stronger brain–behaviour relationships also generalize more equitably across populations, or whether consistent performance gaps between ethnic/racial groups persist regardless of overall predictive accuracy.

Third, potential strategies for mitigating these disparities remain largely untested. Multimodal stacking approaches [21,23] have been shown to reliably enhance predictive accuracy for traits such as cognitive functioning by integrating information across multiple neuroimaging phenotypes [21,24–26]. However, it remains unclear whether such integration mitigates ethnic/racial bias. Furthermore, the extent of representation required from underrepresented groups to minimize these disparities is unknown, as is whether synthetically changing the number of training data from underrepresented groups through oversampling yields additional improvements.

Here, we address these gaps by systematically benchmarking cross-ethnicity generalizability across a broad array of MRI-derived neuroimaging phenotypes in predicting cognitive functioning. We trained predictive models of cognitive functioning on 80 unimodal and 11 multimodal neuroimaging phenotypes. For each neuroimaging phenotype, we compared four training strategies: models trained on (1) All available data (majority White American), (2) Randomly Selected White Americans only (RandWA-only), matched in size to the available African American sample, (3) African Americans only (AA-only), and (4) Balanced African American + Randomly Selected White American subsamples (Balanced AA+RandWA). To assess fairness, we quantified ethnicity-specific prediction errors by calculating the mean absolute error (MAE) separately for White American (WA) and African American (AA) participants in the test sets. A higher MAE in a test group indicated poorer prediction performance for that group. This design allowed us to directly compare performance disparities between groups across models and neuroimaging phenotypes. We derived an Ethnicity Bias Index based on the difference in MAE between WA and AA participants for models trained on RandWA-only and AA-only data, and tested whether neuroimaging phenotypes with higher predictive power exhibited reduced bias. Finally, we extended the analysis to multimodal stacked models to evaluate whether integration of multiple neuroimaging phenotypes improves both accuracy and fairness. In addition to benchmarking existing training strategies, we also explored how incremental increases in AA representation influence prediction accuracy and whether synthetically increasing the number of AA participants through oversampling helps.

Our overarching aim was to provide the first systematic, modality-wide evaluation of ethnicity-related bias in predictive neuroimaging models of cognitive functioning and to test whether boosting predictive accuracy—through feature selection, multimodal integration, or synthetic oversampling—reduces or perpetuates disparities.

## Results

To evaluate ethnicity-related bias, we analysed data from the ABCD dataset, focusing on White and African American participants in groups matched across demographic and socioeconomic factors [27] (Figure 1A). Cognitive functioning was measured using the NIH Toolbox total score [28,29]. We derived 80 unimodal neuroimaging phenotypes from structural MRI (sMRI), diffusion tensor imaging (DTI), functional MRI (fMRI) task contrasts, and functional connectivity (FC), plus 11 multimodal stacked combinations. Models were trained under four strategies: All (majority WA), RandWA-only, AA-only, and Balanced AA+RandWA (Figure 1B). Performance was assessed via mean absolute error (MAE; lower values indicate better predictions), with disparities quantified using an Ethnicity Bias Index (difference in MAE gaps between RandWA-only and AA-only models; lower absolute values indicate less bias). We also tested the effect of incremental and oversampling of AA on model predictions (Figure 1C). Full details are in the Methods and Supplementary Information.

**Figure 1.**
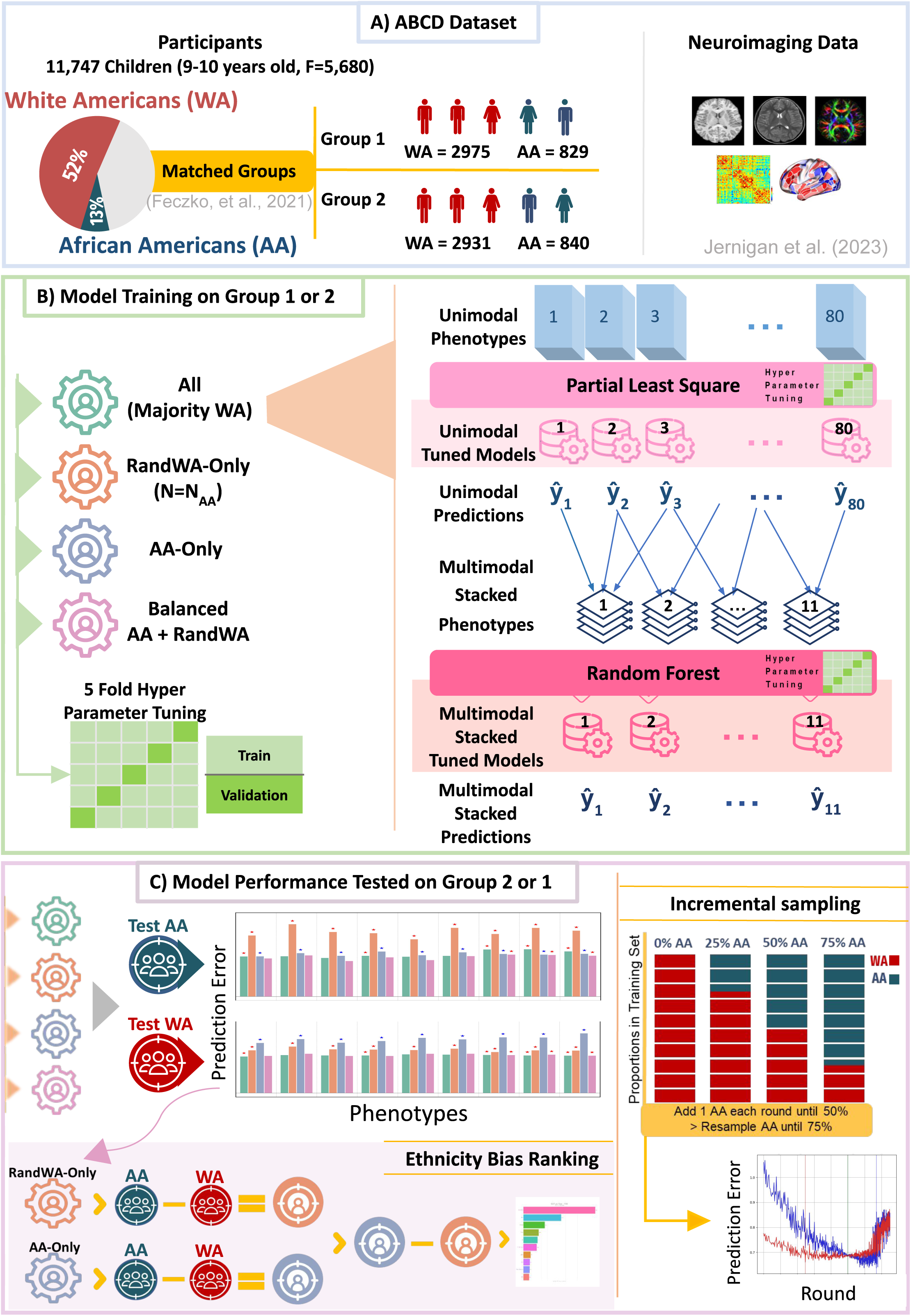
Overall design and information about participants. **(A)** Matched groups and data extracted from the ABCD. **(B)** The pipeline for building the cognitive prediction models and training strategies. **(C)** Evaluation of prediction error across different neuroimaging phenotypes and training strategies in the test set.

### Prediction error across modalities and training strategies

#### sMRI and DTI

For sMRI phenotypes, the RandWA-only models performed significantly better on WA test participants, as indicated by lower MAE compared to AA test participants, based on permutation tests. The All models, trained on the full dataset with a majority WA participants, achieved overall higher accuracy than the RandWA-only model but continued to perform significantly better for WA than AA participants. The AA-only models showed the opposite trend, performing significantly better for AA than WA participants. The Balanced models had better accuracy relative to the ethnicity-specific models and performed on par to the All model but still yielded significantly lower MAE for WA than for AA participants (Figure 2A). See Supplementary Figures 2A and 4A for coefficient of determination (*R2*) and Pearson correlation (*r*) across training strategies.

**Figure 2.**
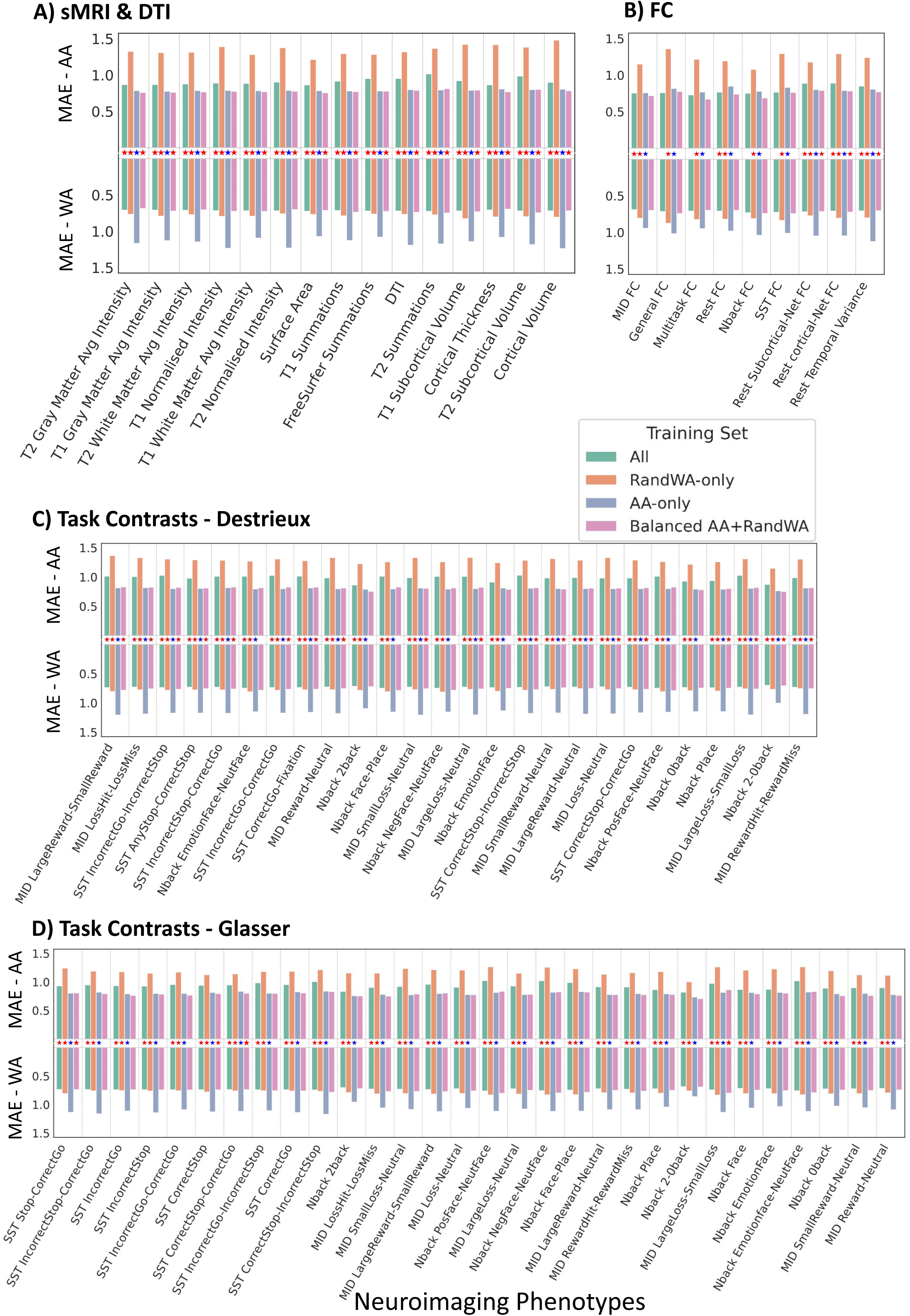
Average prediction error (MAE) for test AA and WA participants across different neuroimaging phenotypes: **(A)** sMRI and DTI, **(B)** FC, **(C)** Task contrasts parcellated with Destrieux [31], and **(D)** Task contrasts parcellated with Glasser atlas [30]. For a given model, higher MAE reflects worse predictive performance. Blue stars indicate significantly better performance in tests AA (i.e. lower MAE) and red stars indicate significantly better performance (i.e. lower MAE) in test WA based on the permutation tests (p-value < 0.05).

#### Functional Connectivity (FC)

For FC phenotypes, RandWA-only models predicted WA participants more accurately, while AA-only models predicted AA participants more accurately. For task-related functional connectivity (Nback, SST, Multitask and General FC), the All and Balanced models achieved similar MAE across WA and AA test groups with no significant difference revealed. Their prediction errors for AA were numerically similar to their AA-only counterparts. Among resting-state phenotypes, this pattern of similar performance across AA and WA participants only held for the balanced model of rest FC (Figure 2B). See Supplementary Figures 2B and 4B for coefficient of determination (*R2*) and Pearson correlation (*r*) across training strategies.

#### Task-based fMRI contrasts

For task-based contrasts, the RandWA-only models performed significantly better on WA test participants, as indicated by lower MAE compared to AA participants. The All model reduced overall errors but continued to perform significantly better for WA than AA participants. In contrast, the AA-only model performed significantly better for AA participants. For contrast phenotypes parcellated with the Glasser atlas [30], the Balanced model achieved comparable MAEs across WA and AA groups, showing no significant performance differences for most task contrasts (26 out of 30) (Figure 2C). However, when using the Destrieux atlas [31], this pattern of comparable performance was observed in only seven out of 26 contrasts (Figure 2D). See Supplementary Figures 2C&D and 4 C&D for coefficient of determination (*R2*) and Pearson correlation (*r*) across training strategies.

#### Multimodal models

Multimodal models were implemented as stacked models that integrated multiple neuroimaging phenotypes. These models exhibited a pattern similar to ethnicity-specific models, with significantly lower prediction error (i.e., better performance) within the ethnic group on which they were trained. When applying the All model, MAE remained significantly lower for WA than AA participants (Figure 3). The Balanced model showed a comparable pattern: only five multimodal models, those combining predictions from task-specific neuroimaging phenotypes, demonstrated non-significant differences between AA and WA. See Supplementary Figures 3 and 5 for coefficient of determination (*R2*) and Pearson correlation (*r*) across training strategies.

**Figure 3.**
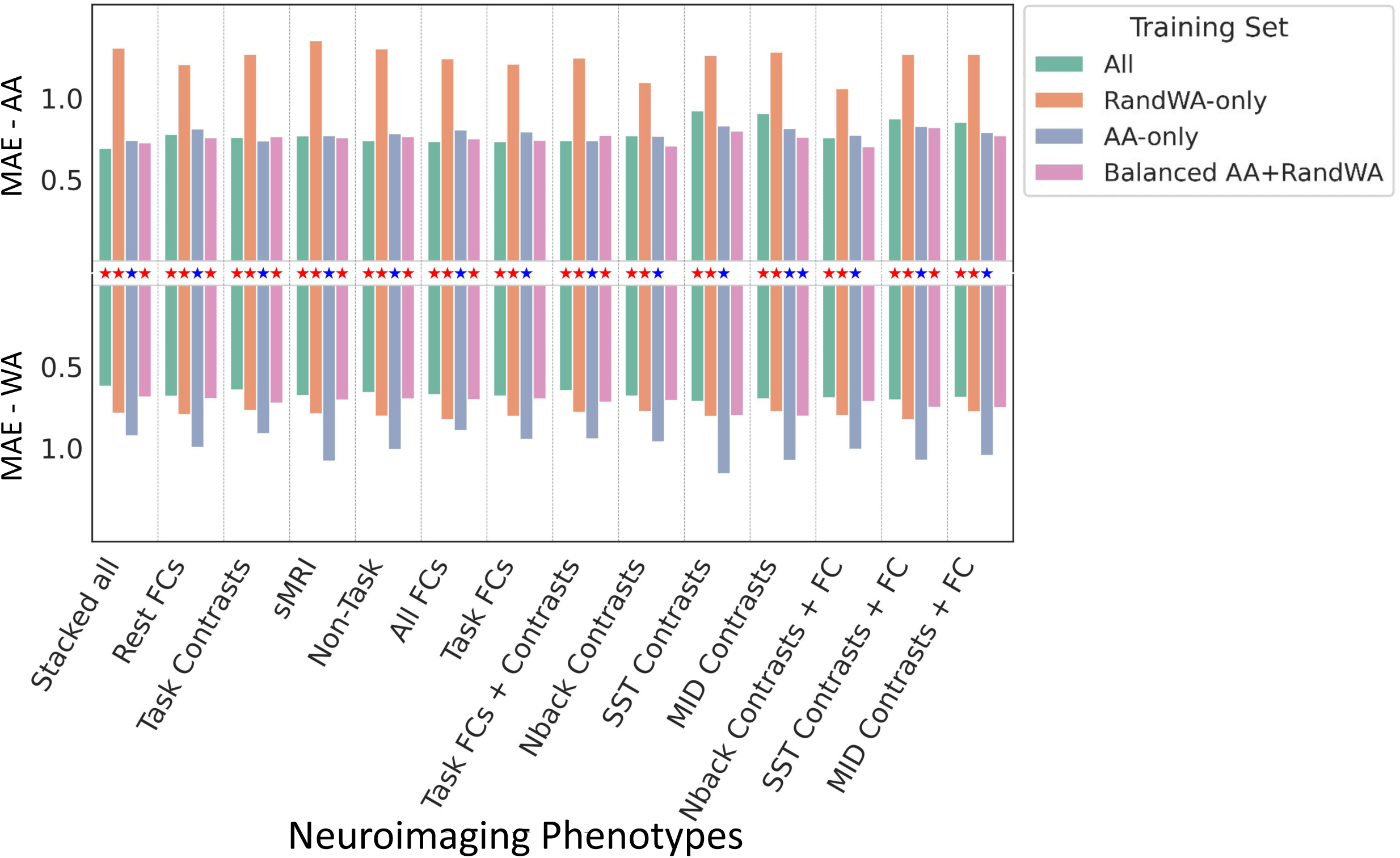
Average prediction error for test AA and WA participants across multimodal stacked neuroimaging phenotypes. For a given model, higher MAE reflects worse predictive performance. Blue stars indicate significantly better performance in tests AA (i.e. lower MAE) and red stars indicate significantly better performance (i.e. lower MAE) in test WA based on the permutation tests (p-value < 0.05).

### Ethnicity Bias Index

The Ethnicity Bias Index was derived for each of the 80 unimodal and 11 multimodal neuroimaging phenotypes by subtracting the group-specific MAE differences observed in RandWA-only and AA-only models. Smaller absolute Ethnicity Bias Index values indicated less ethnicity-related bias. As shown in Figure 4A, ranking neuroimaging phenotypes by Ethnicity Bias Index showed that functional measures, particularly Nback 2-0back contrast and task and rest functional connectivity were among the least biased. Structural MRI phenotypes consistently showed the largest Ethnicity Bias Index values, indicating greater disparities. A similar pattern was seen when ranking multimodal neuroimaging phenotypes: models combining predictions from Nback-related neuroimaging phenotypes and different functional connectivity showed lowest bias and combinations of different sMRI showed the highest. The least biased multimodal phenotype had a higher bias score compared to its unimodal counterpart. The most biased multimodal phenotype had a smaller bias score than its unimodal counterpart (Figure 4B).

**Figure 4.**
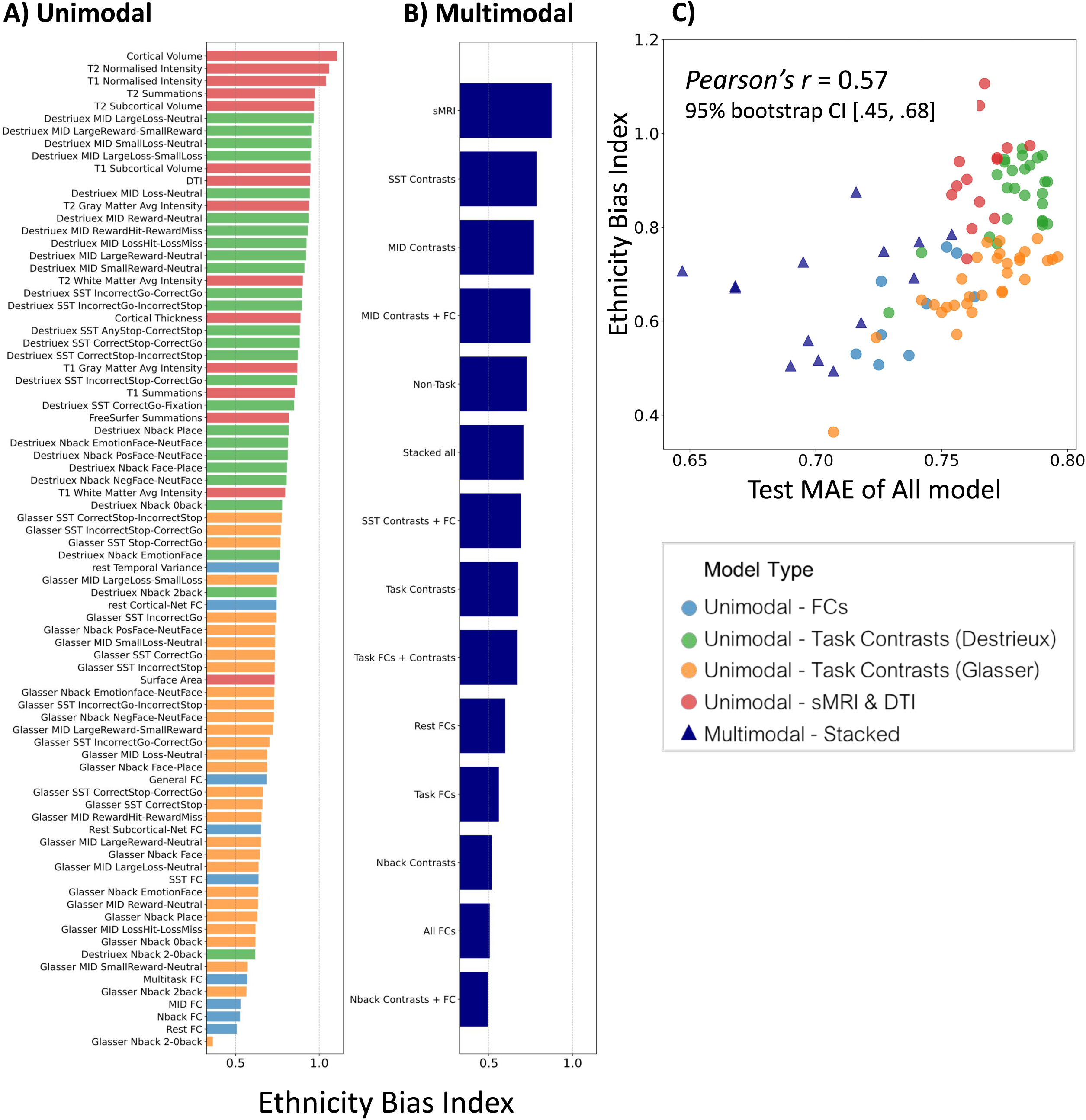
Ranking of **(A)** Unimodal and **(B)** Multimodal neuroimaging phenotypes based on Ethnicity Bias Index. **(C)** The correlation between test MAE of the model trained on all participants and the ethnicity bias index score across both unimodal and multimodal neuroimaging phenotypes. A lower ethnicity bias score for a phenotype indicates reduced favouritism toward any one ethnic group over the other.

### Relation between performance and bias

Across all unimodal and multimodal neuroimaging phenotypes, lower predictive error—calculated by applying the All model on the whole test set, was associated with lower Ethnicity Bias Index score (*r*(89) = .57, 95% bootstrap CI [.45, .68], *p* = 0.0000, Figure 4C). For unimodal phenotypes, the correlation between the All model performance on the whole test set and the corresponding Ethnicity Bias Index values was *r*(78) = .62, 95% bootstrap CI [.44, .75], *p* = 0.0000, indicating that models yielding stronger predictions tended to also yield smaller disparities between AA and WA.

Multimodal stacked models, integrating predictions across multiple neuroimaging phenotypes, achieved lower predictive error than unimodal models. However, bias values in multimodal models were not improved beyond unimodal neuroimaging phenotypes and rather were in the mid-range (Figure 4C). The association between All model performance (MAE) and Ethnicity Bias Index was weaker, with a correlation of *r*(9) =.3, 95% bootstrap CI [-.1, .64], *p* = .3.

To examine how feature contributions to predictions vary with the ethnicity of the training sample, we computed, for each unimodal phenotype, the difference in Partial Least Square (PLS) regression coefficients between models trained on WA-only and AA-only samples. These coefficient differences quantify how the contribution of each brain region or network shifts when models are trained on AA versus WA samples. In Figure 5, we mapped these feature weights back onto brain space for the ten least and most biased unimodal phenotypes.

**Figure 5.**
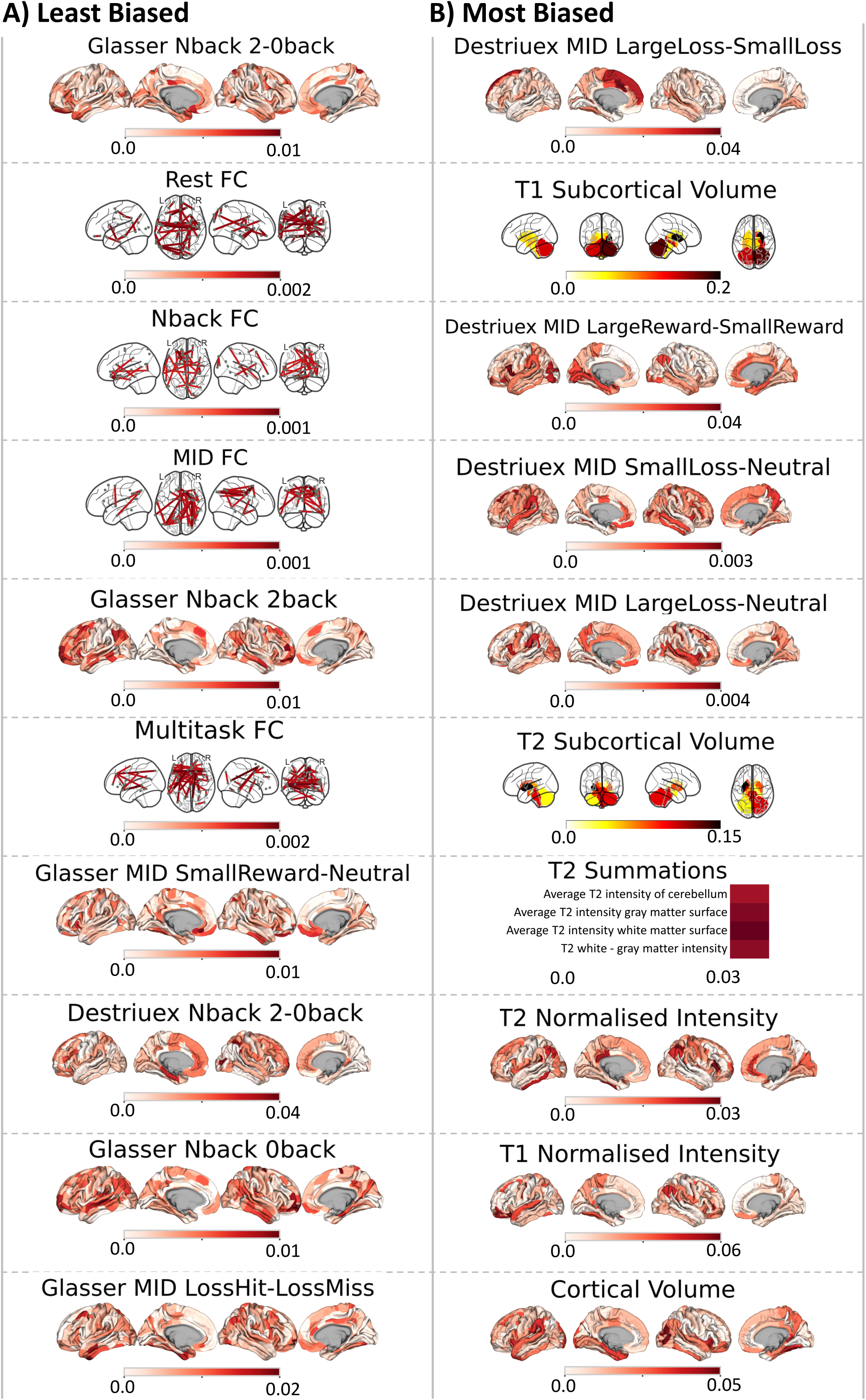
The difference in PLS coefficients when training sample changes from WA-only to AA-only for the ten **(A)** least biased and **(B)** most biased unimodal phenotypes.

### Effects of incremental aa representation and synthetical oversampling

The incremental sampling analysis revealed that increasing the proportion of AA participants in the training set reduced MAE for AA test participants up to approximately the 50% balance point. Beyond this threshold, additional AA representation through synthetical oversampling did not further improve AA performance; in some cases, performance plateaued or worsened. For sMRI phenotypes, a slight decrease in AA test prediction error was observed with additional AA sampling, but this was accompanied by increased error for WA test participants. For task-based and functional connectivity phenotypes, increasing AA representation beyond 50% did not yield further benefits and in some cases increased error (Figure 6).

**Figure 6.**
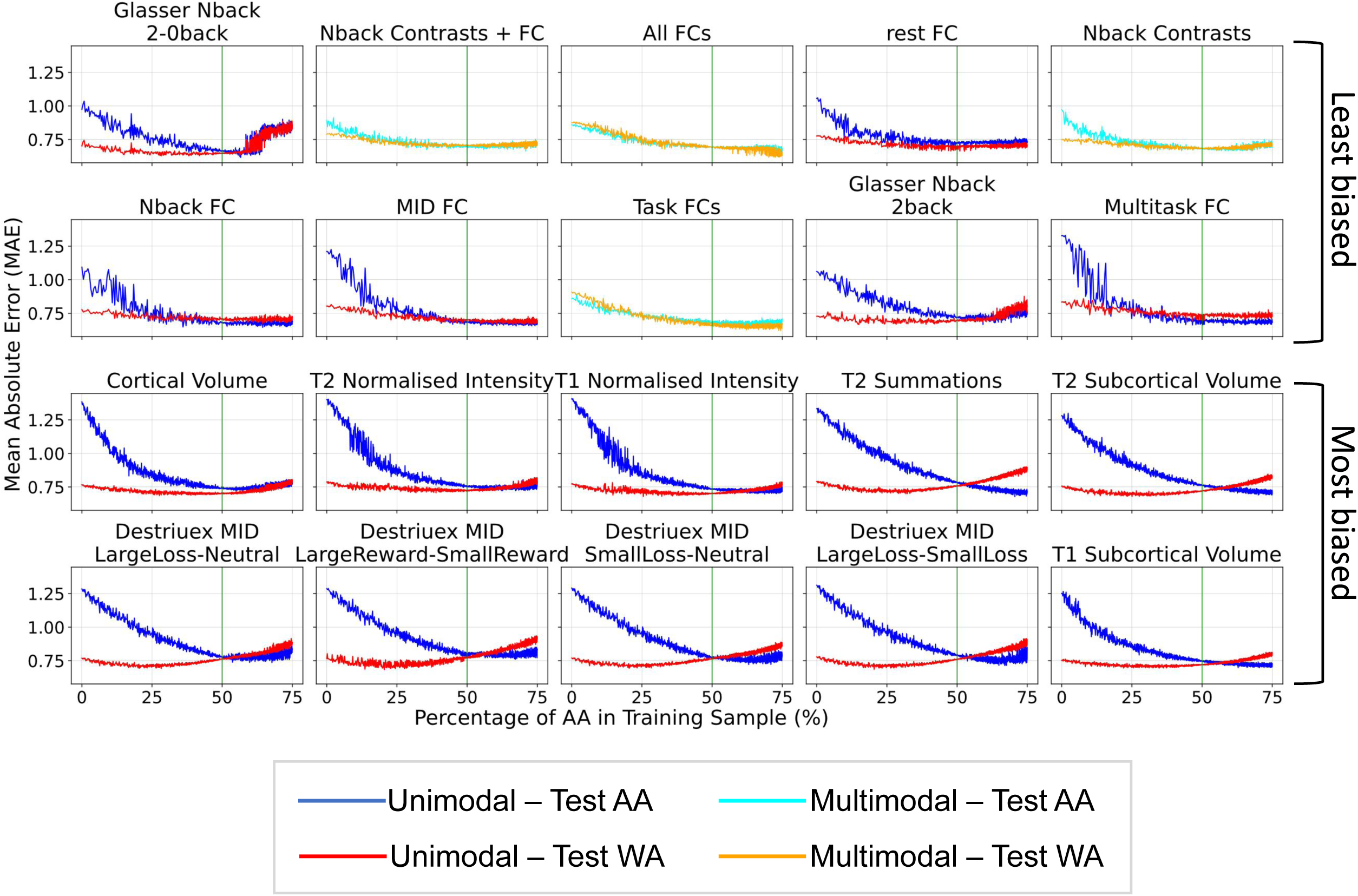
The effect of incremental addition of AA to training set on average prediction errors in test AA and WA for the ten least and most biased among unimodal and multimodal neuroimaging phenotypes. The green vertical line indicates balanced training sample including same number of AA and WA participants. A lower MAE reflects improved predictive performance.

## Discussion

This study provides a systematic benchmark of ethnicity-related generalization in MRI-based predictive modelling of cognition. Several key findings emerged:

### MRI Neuroimaging phenotypes differ in sensitivity to training composition and generalizability

Across all MRI modalities, models trained on a single ethnic group achieved their best performance when applied to that same group. The All models—trained on participants sampled without regard to ethnicity and therefore dominated by White participants—performed better on White participants, consistent with prior findings for resting-state fMRI [18]. Together, these results suggest that the brain–cognition associations captured by predictive models are systematically shaped by the racial/ethnic composition of the training sample.

Balanced AA+RandWA training showed promise in reducing performance disparities between AA and WA test participants, notably without compromising performance for WA participants—a common limitation of fairness optimization methods [32,33]. However, with structural MRI phenotypes, prediction accuracy for AA participants still lagged behind that of WA participants. In contrast, task-based fMRI phenotypes, such as Nback 2–0back and 2back contrasts, were more equitable, with balanced training often eliminating group differences. Task functional connectivity was least sensitive to the ethnic composition of the training set, as both the Balanced and All (majority-White) models performed similarly for AA test participants. Overall, these results highlight that fairness in predictive performance is strongly shaped by both imaging modality and the composition of the training data.

The Ethnicity Bias Index provided a systematic way to rank MRI neuroimaging phenotypes. Task-based contrasts and functional connectivity consistently fell among the least biased neuroimaging phenotypes, while sMRI measures were the most biased. Importantly, within a modality, neuroimaging phenotypes varied: for example, Nback contrasts (0back, 2back and 2-0back) which manipulated working memory load were less biased than other task contrasts.

### Predictive strength is linked to lower bias

Neuroimaging phenotypes with higher predictive accuracy tended to show lower bias, with a moderate correlation between MAE and the Ethnicity Bias Index. This suggests that, although ethnic/racial bias was not fully eliminated, robust brain–behaviour associations generalized more fairly across populations and at least some disparities reflect model fragility rather than intrinsic group differences.

### Predictive gains do not mitigate ethnic/racial bias

The stacked-all model, which combined predictions from all neuroimaging phenotypes, achieved higher predictive performance than any unimodal model when trained on all participants. More broadly, most multimodal models outperformed unimodal models in terms of predictive accuracy across the test set. However, these accuracy gains did not translate into greater fairness. Ethnicity-specific multimodal models favoured their own group, while All and Balanced multimodal models favoured WA participants. Across feature combinations, multimodal models showed intermediate Ethnicity Bias Index values, indicating that their fairness largely depended on the properties of the constituent features. Thus, while multimodal stacking boosted predictive performance, it did not resolve disparities, paralleling lessons from genomics that gains in predictive power do not guarantee equitable outcomes [34].

### Balanced sampling as the fairness ceiling

The incremental sampling analysis provides additional insight into the relationship between dataset composition and bias. As expected, increasing AA representation improved performance for AA test participants up to a balanced training distribution. However, adding AA participants beyond this balance point—via repeated sampling—did not produce further gains. For sMRI, oversampling improved AA predictions slightly but reduced WA performance, suggesting models may be learning group-specific structural patterns in opposing directions. These findings suggest diminishing returns to oversampling beyond balanced representation as a potential solution. Across all 80 unimodal and 11 multimodal phenotypes, no strategy achieved lower ethnicity-specific prediction error than balanced sampling. This includes models trained on the full imbalanced dataset, ethnicity-specific models, multimodal stacked ensembles, and synthetic oversampling. The superiority of balanced sampling stems from its preservation of the full target distribution for the underrepresented group, eliminating extrapolation error without sacrificing exposure to the majority distribution. In the absence of new data collection, balanced subsampling is the optimal mitigation strategy for ethnic bias in neuroimaging-based cognitive prediction.

### Factors contributing to the differences in prediction performance

These results extend prior literature [18] by showing that ethnicity-related bias is modality-dependent. A consistent trend was that neuroimaging phenotypes with higher predictive strength also tended to exhibit lower bias, suggesting that robust brain–behaviour associations may generalize more equitably across populations, whereas weaker predictors may be more fragile to sample composition. This raises the possibility that methodological improvements at the levels of data collection, pre-processing, and algorithm design could simultaneously enhance both accuracy and fairness. However, our findings also demonstrate that fairness does not automatically follow from improved accuracy—multimodal models increased predictive performance but left disparities largely intact.

The observed modality differences—task-based phenotypes being less sensitive to training composition, while structural phenotypes remained more vulnerable—point toward additional methodological sources of bias. Widely used volumetric and surface templates (e.g., MNI152, fsaverage, fs_LR) were constructed primarily from White/European-descendant samples. Applying these templates to non-European populations may increase deformation or segmentation errors, amplifying disparities in sMRI-based prediction. Prior work has shown that population-specific templates can improve segmentation accuracy and reduce deformations in Chinese cohorts [35–37], yet comparable African-specific templates have not been developed. In contrast, task contrasts and task-regressed functional connectivity reduce reliance on structural alignment by emphasizing condition-specific within-subject differences or residual fluctuations in the time series, thereby attenuating biases introduced by structural templates or atlas boundaries. Further evidence comes from our comparison of parcellation schemes: Glasser parcellation [30] yielded both stronger predictions and markedly lower bias compared to Destrieux [31]. Although both atlases were derived from a predominantly White HCP sample, their design differs in ways that may influence cross-ethnic generalizability. The Glasser atlas integrates multiple modalities, including myelin maps, task-related response, and connectivity, to create functionally homogeneous parcels [30,38]. This multimodal basis may make it less sensitive to ancestry-related morphological variation. In contrast, the Destrieux atlas is based primarily on cortical folding patterns such as sulci and gyri [31], which are known to vary across populations [39–41]. This reliance on anatomical landmarks may explain why Destrieux-based phenotypes exhibited larger bias despite only modest differences in predictive performance. Further supporting this view: for Glasser-based contrast and functional connectivity neuroimaging phenotypes, a balanced training sample largely eliminated performance differences between AA and WA test groups. This suggests that, although recent work has documented population differences in functional network topography (Yang et al., 2023), such variability may play a smaller role in cognitive prediction than structural factors embedded in pre-processing and atlas choice.

Together, these observations suggest that modality-dependent disparities reflect both the predictive robustness of phenotypes and the ways in which pre-processing pipelines embed structural assumptions.

### Limitations

Our analyses were limited to WA and AA participants in the ABCD dataset and may not generalize to other underrepresented groups. Also, we relied on widely-used pre-processing pipelines and standard templates—such as the MNI152 brain template—which may embed structural biases that disproportionately affect sMRI-based prediction. The MNI152 template was constructed from 152 healthy young adult MRI scans, primarily from Western populations [42]. The Human Connectome Project (HCP) database—which many surface/atlas pipelines draw from—reports that ∼74.7% of its full sample identified as White/Caucasian [30,43].

### Recommendations for future use of predictive models for neuroimaging

From a translational perspective, these findings underscore the risks of deploying predictive neuroimaging models without explicit attention to data composition. Unequal accuracy across groups could reinforce existing health disparities—a concern long recognized in genomics, where polygenic scores trained in European ancestry cohorts show limited transferability to other populations [12,16,44]. Similar principles apply to neuroimaging: expanding dataset diversity and adopting balanced training designs are essential for equitable generalization, particularly since balanced training sample as a bias-mitigation technique did not exhibit the accuracy–fairness trade-off cautioned in fairness studies [32,33]. Balanced sampling thus represents the practical upper bound for equitable prediction — a simple, cost-free strategy that outperforms complex multimodal integration and data augmentation.

Modality choice also matters. Selecting neuroimaging phenotypes less sensitive to pre-processing biases—such as task-based contrasts or task-regressed functional connectivity—may improve fairness. Therefore, when predictive models are applied as research or clinical tools, it may be more appropriate to prioritise such phenotypes that demonstrate lower sensitivity to training-sample composition.

Developing ethnicity-specific models, analogous to subgroup-specific polygenic scores, might improve within-group predictions but risks institutionalizing segregated tools and diverting attention from inclusivity. If models trained on large White-majority datasets cannot generalize, this reflects deeper representational limits in how current neuroimaging frameworks capture human diversity. The priority should be developing processing pipelines and models that are both accurate and demographically inclusive.

Algorithmic fairness interventions offer complementary solutions. These techniques aim to reduce model dependence on sensitive attributes such as ethnicity and improve generalization across population by removing information about group membership from model representations (adversarial debiasing), aligning data distributions across group (domain adaptation) or adjusting sample importance during training (reweighting) [10,32,45,46]. Beyond algorithmic solutions, advances in more inclusive brain templates and functionally representative atlases could further mitigate structural biases embedded in current pre-processing standards. Together, these efforts represent necessary steps toward equitable and generalizable neuroscience.

## Conclusion

In sum, this study provides the first modality-wide benchmark of ethnicity-related bias in predictive neuroimaging. Disparities varied across modalities and model types, with structural phenotypes consistently more biased and task-based and connectivity phenotypes more equitable. Predictive strength was moderately associated with fairness, but stacking did not resolve disparities. Balanced training emerged as the upper bound for equitable predictions, but oversampling beyond balance yielded diminishing returns. These results underscore that ensuring equitable neuroimaging-based prediction will require diverse datasets, careful feature selection, and explicit bias-mitigation strategies. As neuroimaging-based prediction advances toward clinical translation, addressing bias is both a scientific necessity and an ethical imperative to ensure that progress in precision neuroscience benefits all populations.

## Methods

### Participants

Data were drawn from the Adolescent Brain Cognitive Development (ABCD) Study Curated Annual Release 5.1 (DOI:10.15154/z563-zd24), a longitudinal neuroimaging project with a diverse U.S. cohort [47]. The baseline cohort included 11,878 participants (48% female; 52% male) aged 9–10 years. To examine ethnicity-related generalization, we selected WA and AA participants from the two bigger matched groups of the ABCD-BIDS Community Collection (ABCC-2.0) [48], where participants were matched across 21 sites based on a combination of clinical (history of anaesthesia), demographic (age, sex, race/ethnicity) and socioeconomic variables (family income, family type, household size) to ensure comparability between groups [27]. For the purpose of current study, we interchangeably used the word race and ethnic to refer to the characterisation of WA and AA groups.

### Training strategies

We created four ethnicity-based training strategies to evaluate the effect of ethnicity on model performance. The “All model” included all (majority White) participants in the training matched group. The “RandWA-only model” trained on White American participants, randomly selecting a number of White Americans equal to the number of African American participants to ensure comparability. The “AA-only model” trained exclusively on African American participants. The “Balanced model” included all African American participants along with an equal number of randomly selected White Americans, creating an ethnically balanced training set.

The overall study design, including the cross-validated training–testing framework, the ethnicity-based training strategies, and the corresponding sample sizes for each group, is illustrated in Figure 1.

### Cognitive measures

The cognitive phenotype of interest was the total cognitive composite score from the NIH Toolbox [28,29]. We selected the total cognition composite as the primary outcome because it exhibits higher test–retest reliability and reduced measurement noise relative to individual task scores [49–51], and has been widely used in prior neuroimaging-based prediction studies within ABCD and related developmental cohorts [52–54]. Aggregating across domains yields a more stable and robust phenotype for predictive modelling, mitigates task-specific idiosyncrasies, and improves sensitivity to distributed brain–behaviour relationships that are unlikely to be captured by single cognitive measures [51,55,56]. See Supplementary Information for details of each cognitive test.

### Neuroimaging phenotypes

Neuroimaging data were drawn from the Adolescent Brain Cognitive Development (ABCD) Study, integrating two complementary sources: (i) post-processed tabulated summary measures released by the ABCD consortium (version 5.1, DOI:10.15154/z563-zd24) and (ii) additional phenotypes derived from custom processing of minimally pre-processed data from the ABCD-BIDS Community Collection (ABCC) [48]. The ABCD tabulated datasets provide quality-controlled regional measures and task contrasts parcellated using the Destrieux cortical atlas [31] and FreeSurfer [57] subcortical segmentation. To extend spatial resolution and connectivity and structural coverage beyond these summaries, we further processed ABCC data using outputs from fMRIPrep (v20.2.0) and the Human Connectome Project (HCP) minimal pre-processing pipeline. See Hagler et al. [58] and Supplementary Information for more details about how we processed the neuroimaging data.

After applying ABCD-recommended quality control and exclusion procedures—including removal of Site 22 due to limited sample size, correction of site ID errors, exclusion of participants with reported vision problems, scan and task quality flags—the final neuroimaging sample sizes ranged from approximately 4,000 to 11,177 participants, depending on modality and phenotype. Importantly, exclusions were applied at the phenotype level rather than via listwise deletion, allowing participants to contribute data to phenotypes that met quality criteria. In total, this pipeline yielded 80 distinct neuroimaging sets (summarized in Supplementary Information, Supplementary Figure 1).

#### Functional MRI (fMRI) Task Contrasts: 56 sets

Task-evoked fMRI BOLD response were derived from three ABCD tasks: the Emotional N-back task (working memory and affective processing) [59], the Monetary Incentive Delay (MID) task (reward and punishment processing) [60], and the Stop Signal Task (SST; inhibitory control) [61]. From the ABCD tabulated data, we used 26 precomputed contrasts (9 N-back, 10 MID, 7 SST), parcellated with the Destrieux cortical [31] and FreeSurfer subcortical [57] atlases [58].

To complement these measures, we computed an additional 30 task contrasts (10 per task) from ABCC data using a first-level General Linear Model (GLM) framework implemented primarily in Nilearn [62]. Task event files were converted from E-Prime format to BIDS-compatible TSV files, with event onsets aligned relative to post-calibration task start times. Pre-processing incorporated nuisance regression informed by prior recommendations, including cosine drift regressors, ten anatomical CompCor components, and six rigid-body motion parameters with their first temporal derivatives. Non-steady-state volumes were removed in a scanner-specific manner prior to modelling.

GLMs were estimated using ordinary least squares with the SPM canonical hemodynamic response function. For each contrast of interest, beta coefficients were estimated at the run level and combined across runs using variance-weighted fixed effects. Contrasts were designed to isolate specific cognitive processes, such as working memory load (e.g., 2back, 0back, 2-0back), reward magnitude (e.g., large reward vs. neutral), response type (e.g., hit vs. miss), and inhibitory control (e.g., successful stop vs. go). Final contrast maps were parcellated into the 379-region Glasser–FreeSurfer [30,57] atlas to produce regional effect sizes.

#### Functional MRI Connectivity (fMRI FC): 9 sets

Functional connectivity phenotypes were derived from both ABCD tabulated data and ABCC-based analyses. From the ABCD releases, three resting-state FC sets were included: (i) temporal variance measures reflecting amplitude of low-frequency fluctuations across 333 Gordon cortical parcels [63] and 19 subcortical regions [57] ; (ii) subcortical-to-network connectivity, defined as average correlations between each subcortical region and 13 large-scale cortical networks; and (iii) within- and between-network cortical connectivity across these networks. Using ABCC data, we additionally computed region-to-region FC matrices at the 379-region resolution Glasser–FreeSurfer [30,57] for resting state and each of the three tasks, yielding four further sets. Nuisance regression and dummy-scan removal followed the same procedures as for task activation analyses. Motion outliers were identified using framewise displacement (>0.5) and DVARS (>1.5), with flagged volumes incorporated as regressors. Participants with more than have flagged volumes were excluded. Given the multiband acquisition and developmental sample, a relatively lenient motion threshold was adopted to balance artifact mitigation with signal preservation. Residual time series were extracted after regressing out nuisance variables and, for task data, task-evoked activity. Extreme values were attenuated via de-spiking prior to connectivity estimation. Residuals were concatenated across runs and parcellated into the 379 regions. Functional connectivity was computed using Pearson correlation, followed by Fisher z-transformation and vectorization. In addition to task-specific and resting-state FC, we derived multitask FC by concatenating residuals across all three tasks and a general FC measure by concatenating residuals across rest and all tasks, consistent with evidence that aggregated data improve reliability and behavioural prediction [64–66] (see Supplementary Information).

#### Structural MRI (sMRI) and Diffusion Tensor Imaging (DTI): 15 sets

Structural MRI phenotypes were derived primarily from FreeSurfer [57] processing of T1- and T2-weighted images. We included 14 structural phenotypes capturing complementary aspects of brain morphology and tissue properties, comprising regional measures of cortical thickness, surface area, volume, and subcortical volumes, parcellated using the Destrieux cortical atlas [31] and FreeSurfer subcortical segmentation [57]. In addition to morphometric measures, we incorporated gray- and white-matter intensity which reflect local signal properties sampled at cortical surfaces or within subcortical masks. Corresponding normalized intensity measures were also included to reduce scanner- and site-related variability. Beyond these region-specific metrics, we leveraged additional T1- and T2-weighted intensity–based phenotypes provided in the ABCD tabulated datasets. These phenotypes summarize average signal intensities across cortical and subcortical regions, as well as global tissue classes, and include whole-brain “summation” measures that aggregate intensity information across gray matter, white matter, and cerebellar regions, along with contrasts between tissue types (e.g., gray–white matter differences). Compared with gray-and white-matter intensity measures derived directly from FreeSurfer, which index localized tissue signal at specific anatomical surfaces, the T1- and T2-based summaries are designed to capture broader regional and global signal characteristics sensitive to tissue composition, including myelination, water content, and other microstructural properties [67,68]. We additionally included a global FreeSurfer summations comprising five whole-brain metrics: estimated intracranial volume, total cortical gray matter volume, total cortical white matter volume, total subcortical gray matter volume, and the ratio of brain segmentation volume to intracranial volume. Together, these diverse structural representations were selected to capture both localized and global aspects of brain structure and tissue composition, providing complementary information relevant to individual differences in cognitive development.

The Diffusion phenotype consisted of fractional anisotropy (FA) values extracted from 23 major white-matter tracts identified using AtlasTrack. FA provides an index of directional water diffusion and was included to capture complementary aspects of white-matter microstructural organization relevant to cognitive development [69].

### Predictive modelling

Models were trained on one matched group [27] and tested on the other, ensuring that test participants were entirely independent of the training set.

All neuroimaging phenotypes and the cognitive scores were standardized (z-scored) based on the training data, and the same standardization parameters were applied to the test set. Hyperparameters were optimized using a five-fold cross-validation within each training set, with negative mean squared error (NMSE) as the evaluation metric.

Prediction models were developed in two frameworks:

#### Unimodal models (80 models total)

For each of the 80 neuroimaging phenotypes (derived from task-based fMRI, resting-state fMRI, sMRI, and DTI), we trained a separate predictive model using Partial Least Squares (PLS) regression, a supervised dimensionality reduction approach that maximizes covariance between neuroimaging features and the cognitive target [70]. The number of PLS components was tuned via grid search (30 components maximum; for phenotypes with fewer than 30 variables, all possible components were considered). PLS was selected not only for its ability to handle high-dimensional and highly collinear data (e.g., functional connectivity matrices), but also because prior analyses in our primary work demonstrated comparable predictive performance between PLS and Elastic Net regression in this dataset, and independent benchmarking study has shown Elastic Net to perform well in neuroimaging-based cognitive prediction [20]. Given this equivalence in performance, PLS was preferred here due to its explicit dimensionality reduction, which yields compact latent representations while retaining variance most relevant to the cognitive outcome.

#### Multimodal (stacked) models (11 configurations)

To integrate information across phenotypes and capture potential non-linear interactions, we trained 11 multimodal models using late fusion [71]: predictions from the relevant unimodal PLS models were combined as inputs (rather than raw feature concatenation). Each configuration integrated predictions from related groups of unimodal models (e.g., “stacked all” used predictions from all 80 phenotypes; “stacked task contrasts” used predictions from task-based fMRI contrast phenotypes such as N-back, MID, and SST; see Supplementary Figure 1 for full list). The multimodal regressor was a Random Forest [72] with 1,000 trees. Hyperparameters were tuned via inner-fold grid search: maximum tree depth (None or 3–10), and number of features considered at each split (sqrt or log2 of total input features). Random Forest was selected for the multimodal layer due to its robustness to mixed-scale inputs, ability to model non-linear relationships and interactions between modalities, and native handling of missing values tested in previous studies [20,21,73]

We trained 368 models in total, spanning 91 neuroimaging phenotypes (80 unimodal, 11 multimodal) across 4 training strategies (All, RandWA-only, AA-only, Balanced). See Supplementary Figure 1 for a full list of unimodal and multimodal models.

### Model performance

Predicted values from test sets were concatenated across matched groups, and model performance was evaluated using MAE, R2 and Pearson’s r for each training strategy and phenotype. For each model, to assess differences in predictive performance between AA and WA participants in the test set, we used a permutation test. Predicted scores were shuffled 1,000 times to generate a null distribution of performance differences between the two groups. The observed difference was then compared against this null distribution to calculate a p-value. Here, we emphasize MAE as the primary metric for group-wise model comparison because it provides a direct and scale-preserving measure of prediction error that is readily interpretable at the individual level. MAE reflects absolute deviations between predicted and observed values and aligns with common practices in model comparison when assessing performance equity [74]. See Supplementary Information and Supplementary Figures 2-5 for the other metrics.

### Ethnicity Bias Index

To assess ethnicity-specific prediction errors, we computed the difference in MAE between AA and WA in the test set when using the model trained on White Americans (RandWA-only). The same was done for the AA-only model. The Ethnicity Bias Index was then obtained by subtracting the difference observed in the RandWA-only model from the difference observed in the AA-only model. Lower absolute Ethnicity Bias Index values indicate that model predictions are less influenced by participant ethnicity, reflecting less bias. To investigate the relationship between overall predictive performance and bias, we then calculated Pearson’s correlation between Ethnicity Bias Index scores and the All model performance on the whole test set for each unimodal and multimodal phenotype. Confidence intervals were estimated using nonparametric bootstrap resampling (1,000 iterations).

### Incremental sampling and synthetic oversampling

To assess the effect of training sample composition on prediction accuracy, we implemented an incremental sampling procedure. First, we randomly selected a subset of WA participants equal in size to the available AA participants in the training set. Models were trained on this WA set, and MAEs were calculated for WA and AA participants in the independent test set. We then progressively increased the proportion of AA participants by randomly adding one AA participant at a time to the training set, repeating the process until AA participants comprised 75% of the training set. Beyond the 50:50 balance point, we conducted synthetic oversampling. AA participants were randomly re-sampled with replacement to increase their proportion. For each step, the model was retrained, and prediction errors for both the WA and AA test groups were recorded. For the stacked stage, unimodal predictions derived from the Balanced AA+RandWA strategy were used to create stacked neuroimaging phenotypes.

## Acknowledgments

Data used in the preparation of this article were obtained from the Adolescent Brain Cognitive Development (ABCD) Study (https://abcdstudy.org), held in the NIMH Data Archive (NDA). This is a multisite, longitudinal study designed to recruit more than 10,000 children age 9-10 and follow them over 10 years into early adulthood. The ABCD Study® is supported by the National Institutes of Health and additional federal partners under award numbers U01DA041048, U01DA050989, U01DA051016, U01DA041022, U01DA051018, U01DA051037, U01DA050987, U01DA041174, U01DA04110, U01DA041117, U01DA041028, U01DA041134, U01DA050988, U01DA051039, U01DA04115, U01DA041025, U01DA041120, U01DA051038, U01DA041148, U01DA041093, U01DA041089, U24DA041123, U24DA041147. A full list of supporters is available at https://abcdstudy.org/federal-partners.html. A listing of participating sites and a complete listing of the study investigators can be found at https://abcdstudy.org/consortium_members/. ABCD consortium investigators designed and implemented the study and/or provided data but did not necessarily participate in the analysis or writing of this report. This manuscript reflects the views of the authors and may not reflect the opinions or views of the NIH or ABCD consortium investigators. The ABCD data repository grows and changes over time. The ABCD data used in this report came from https://doi.org/10.15154/z563-zd24. DOIs can be found at https://nda.nih.gov/abcd/abcd-annual-releases. The authors acknowledge the use of New Zealand eScience Infrastructure (NeSI) high-performance computing facilities and support services, funded by NeSI collaborator institutions and the Ministry of Business, Innovation & Employment. This research was supported by the New Zealand Health Research Council (Grant numbers: 21/618 and 24/838; awarded to Narun Pat), the University of Otago, and the Neurological Foundation of New Zealand (Grant number: 2350 PRG; awarded to Narun Pat).

## Author contributions

Khakpoor and Pat conceived and designed the study. Data acquisition, analysis and interpretation were conducted by Khakpoor, Pat, Deng and van der Vliet. Khakpoor and Pat drafted the manuscript and performed statistical analyses. All authors critically revised the manuscript. Funding was acquired by Pat. Pat also supervised the project. All authors approved the final version and are accountable for the accuracy and integrity of the work. This manuscript reflects the authors’ views and does not necessarily represent those of any funding body or the ABCD Consortium.

## AI Use Disclosure Statement

During manuscript preparation, the first author used ChatGPT (GPT-5.2, OpenAI) to assist with language editing and refinement. The AI tool was used solely for editorial support. All scientific content, analyses, interpretations, and conclusions were developed by the authors.

## Availability of Data and Materials

ABCD Study data are accessible via https://nda.nih.gov/abcd upon NDA approval. All analysis code is available at https://github.com/HAM-lab-Otago-University/Ethnicity-Benchmark

