## Supplementary Information for "Brain predictive models of cognition fail to generalize across ethnicities: Modality-dependent bias in MRI-based prediction"

### Supplementary Materials

### Cognitive-Functioning

NIH Toolbox composite scores [1] for total cognitive-functioning were derived from seven tasks: Picture Vocabulary measures receptive vocabulary and verbal comprehension [2]. Flanker Inhibitory Control & Attention evaluates attention and response inhibition [3]. List Sorting Working Memory tests working memory and sequencing [4]. Dimensional Change Card Sort assesses cognitive flexibility and set-shifting [3]. Pattern Comparison Processing Speed measures visual processing speed [5]. Picture Sequence Memory evaluates episodic memory and sequential recall [6]. Oral Reading Recognition tests reading ability and word recognition [2]. Together, these tasks provide a comprehensive assessment of cognitive function across multiple domains.

### Neuroimaging:

#### Quality Considerations

Quality control for neuroimaging data followed ABCD-recommended exclusion criteria [7]. Exclusion flags based on metrics such as image quality, MR neurological screenings, and task performance, were assigned to each neuroimaging feature [8]. We did not apply listwise deletion, but rather excluded specific sets of neuroimaging features that failed to meet the criteria. For instance, when participants failed to meet the criteria for Nback task fMRI, but not other sets, we only dropped Nback task fMRI from our analysis. See below under Cognitive Prediction Models on how we dealt with missingness through opportunistic stacking [9]. We also excluded 58 children who had vision problems [10].

ABCD provided post-processed tabulated data with specific atlases. And we use them directly. We then also post-processed the ABCD-BIDS Community Collection (ABCC) data ourselves to complement the post-processed data provided by the ABCD.

### ABCC Pre-processing Pipeline

ABCC is a curated dataset derived from the Adolescent Brain Cognitive Development (ABCD) Study, formatted according to the Brain Imaging Data Structure (BIDS) standard [11]. It provides easy-to-use neuroimaging and behavioral data for researchers. The ABCC includes preprocessed data like MRI images and associated metadata. This collection also includes the outputs of fMRIPrep (version: 20.2.0) and the Human Connectome Project's minimal preprocessing pipelines. We used the generated confounds table from fMRIPrep and the CIFTI file from hcp-pipeline outputs. We converted the text E-prime event files to tsv files using the eprimetotsv.py script by Demidenko and colleagues [12]. The resulting event times are calculated from the task start time after calibration scans. Our first level analysis pipeline was mainly built with Nilearn [13], Nibabel [14] packages .

For the task fMRI in the ABCC, we conducted the following processing steps. First, nuisance regressors were selected and included in the design matrix based on recommendations in the literature [15], [16]. These include a subset of confounds from fMRIPrep to account for potential noise sources: Cosine regressors were added to remove low-frequency noise, such as scanner drift or physiological cycles, which are unrelated to neural activity [15]. Additionally, ten anatomical CompCor components were derived from signals within anatomically defined noise regions, including white matter and cerebrospinal fluid (CSF). These principal components (PC1–PC10), which are unlikely to contain neural activity, were included to account for physiological noise like heartbeat and respiration [17]. Movement regressors, representing head motion parameters calculated during realignment, were also included. These regressors consisted of translations (x, y, z) and rotations (roll, pitch, yaw) in three dimensions, as well as their first derivatives (rate of change). Including these regressors mitigates motion-related artifacts, such as spurious correlations and signal distortions caused by subject movement during the scan [18]. Specific regressors included rot_x, rot_x_derivative1, rot_y, rot_y_derivative1, rot_z, rot_z_derivative1, trans_x, trans_x_derivative1, trans_y, trans_y_derivative1, trans_z, and trans_z_derivative1. To address initial calibration effects, dummy scans were removed from the confounds and image data. The number of removed TRs varied by scanner type: Siemens and Philips scans had 8 TRs removed, GE DV25 scans had 5, and GE DV26 scans had 16 TRs removed [7]. Frame times were calculated based on the number of scans and TR.

The first-level design matrix was created using the SPM HRF model. Subsequently, time series standardization was performed, and a General Linear Model (GLM) analysis was run using Ordinary Least Squares (OLS) as the noise model. Contrasts of interest were defined by creating a contrast matrix. For each contrast, effect sizes were computed by estimating the GLM coefficients. These results were averaged by computing fixed effects, incorporating variance across runs, and calculating the weighted average effect size for contrasts across the two runs. Finally, the averaged effect size maps were parcellated into 379 regions, comprising 360 cortical regions based on the Glasser atlas [19] and 19 FreeSurfer [20] subcortical regions.

For the task and rest fMRI FC in the ABCC, we conducted the following processing steps. We began with the selection of nuisance regressors and the removal of dummy scans, as described earlier. Frame times were calculated based on the number of scans and TRs. Outlier detection and regression were applied by identifying high-motion TRs with framewise displacement (FD) > 0.5 and standardized derivative of root mean square variance over voxels (DVARS) > 1.5, as described in prior literature [21], [22]. FD was calculated as the sum of the absolute displacements in all six motion parameters (translation along x, y, z axes, and rotation about these axes) between successive fMRI volumes. DVARS quantified the root mean square of signal intensity differences across the brain between successive volumes [23]. The identified TRs were flagged as outliers and added as additional nuisance regressors [24]. A censoring mask was created to exclude flagged outliers, their adjacent five TRs, and any time intervals containing fewer than five consecutive sub-threshold TRs. Participants were excluded if their censoring mask flagged more than half of the total timepoints. A lenient FD threshold was used, with no additional motion scrubbing, due to the multiband acquisition, participant demographics, and the importance of preserving information when examining brain-behavior associations [21], [25], [26].

The first-level design matrix was constructed using the SPM HRF model for task fMRI, while no HRF model was applied to rest fMRI. Time series data were standardized, and a de-spiking approach was applied to correct extreme values for connectivity. Specifically, time points exceeding a threshold of 3 standard deviations were replaced with the average value across all time points for each vertex [27]. GLM analysis was performed using OLS as the noise model, and residual signals were extracted after regressing out nuisance variables and, in the case of task fMRI, task-related activity. The residual time series were then concatenated across runs for each participant.

The residual time series were parcellated into 379 regions. Functional connectivity between regions of interest (ROIs) was computed using Pearson's *r*, followed by Fisher's z-transformation of the correlation matrices. These z-transformed matrices were flattened into 1D arrays for subsequent analysis.

#### Functional MRI (fMRI) Tasks

Emotional Nback task combines working memory demands with emotional processing, using stimuli that include emotional faces (positive, negative, and neutral) and places. Participants perform two conditions: a 0-back task, where they identify whether an image matches one shown at the beginning of the block, and a 2-back task, requiring them to determine whether an image matches one presented two trials earlier. This design allows the task to probe neural activity related to working memory, emotion processing, and facial recognition [7], [8].

The Monetary Incentive Delay (MID) task was designed to investigate reward processing. In this task, participants responded to visual stimuli, with their outcomes determined by their reaction time and the specific condition. Depending on the trial, they could win or lose different amounts of money or have no monetary outcome (Neutral). At the end of each trial, feedback indicated whether they won money (Positive Reward Feedback), missed out on winning (Negative Reward Feedback), avoided a loss (Positive Punishment Feedback), or incurred a loss (Negative Punishment Feedback) [7], [28].

The Stop Signal Task (SST) task was designed to study inhibitory control through fMRI by requiring participants to suppress or interrupt their motor response to a ‘Go’ stimulus upon seeing a ‘Stop’ signal. This task probes the neural mechanisms underlying response inhibition and motor control[7], [28].

Using the ABCC, we computed 10 contrasts (see Supplementary figure 1) for each of the fMRI tasks parcellated with Glasser [19] cortical and FreeSurfer[20] subcortical atlases. The contrasts for the Nback task were derived from the design matrix, which represents various task conditions such as stimulus types (places and faces), emotional valences of faces (positive, negative, neutral), and memory loads (two-back and zero-back). These contrasts were constructed to isolate specific effects, including responses to individual stimulus categories (e.g., place or face), combinations of conditions (e.g., emotional faces), and direct comparisons between conditions (e.g., face versus place or positive versus neutral faces). They also allow for the examination of memory load effects by comparing two-back and zero-back conditions. The contrasts for the MID task were derived from the design matrix, which encodes conditions reflecting varying reward and punishment magnitudes (large and small), as well as neutral trials and outcome events (hits and misses). These contrasts were constructed to isolate the neural responses to rewarding versus neutral conditions (e.g., large and small rewards compared to neutral) and similarly for punishment conditions. In addition, contrasts directly compare differences in magnitude (e.g., large versus small rewards or losses) and distinguish between successful versus unsuccessful outcome processing by contrasting hits with misses. The contrasts for the SST task encoded specific task conditions such as stimulus and response type. These contrasts were constructed to isolate neural responses associated with each condition and to directly compare performance differences—such as contrasting CorrectStop with CorrectGo to assess inhibition success, and contrasting IncorrectStop or IncorrectGo with CorrectGo to examine error-related effects. In addition, composite contrasts were created to compare stop versus go trials (e.g., averaging CorrectStop and IncorrectStop relative to CorrectGo) and to differentiate between successful and failed inhibition (e.g., CorrectStop versus IncorrectStop).

The task contrasts available in ABCD tabulated data encompass nine contrasts from Nback, ten from MID and seven from SST parcellated with Destrieux [29] cortical and FreeSurfer [20] subcortical atlases (see Supplementary figure 1).

#### fMRI Functional Connectivity (FC)

In addition to the region-to-region FC during rest and the three tasks in ABCC, we concatenated the residuals from all available tasks to calculate a multitask FC [30] and all available task residuals plus the rest to obtain general FC [31]. The rs-fMRI sets of features in ABCD included temporal variance measurements across 333 cortical [32] and 19 subcortical regions. These values represent the variability over time within each parcellated brain region, capturing the amplitude of low-frequency oscillations [33] ; subcortical-to-network FC (247 features) including the average correlation between each of the 19 subcortical regions and the 13 major cortical networks; and cortical FC (91 features) included the mean correlation values calculated both within and between large-scale cortical networks [7] (see Supplementary figure 1).

#### Structure MRI (sMRI) and Diffusion Tensor Imaging (DTI)

The ABCD collection includes multiple measures extracted from T1-weighted and T2-weighted 3D structural images using FreeSurfer [20]. Cortical and subcortical regions of interest were calculated with Destrieux [29] and FreeSurfer [20] parcellations respectively. We used FreeSurfer's [20] aparc.stats and aseg.stats files to extract three distinct sets of features within ABCC: cortical thickness, surface area, and FreeSurfer summations. The FreeSurfer summations included five features from FreeSurfer [20]: estimated intracranial volume, total cortical gray matter volume, total cortical white matter volume, total subcortical gray matter volume, and the ratio of brain segmentation volume to estimated intracranial volume [34]. We also used 12 other structural sets of features from ABCD tabulated data, including cortical volume, sulcal depth, white-matter averaged intensity, grey-matter averaged intensity, T1 normalized intensity, T1 subcortical averaged intensity, T1 subcortical volume, T1 summations, T2 normalized intensity, T2 subcortical averaged intensity, T2 subcortical volume and T2 summations [7]. Note that T1 summations included average T1 intensity white matter surface, cortical gray matter total area, subcortical gray volume, difference between T1 white and gray matter, cortical gray matter mean thickness, mean sulcal depth, average T1 intensity of cerebellum, average T1 intensity gray matter surface, cortical gray matter total volume. T2 summations included average T2 intensity of cerebellum, average T2 intensity gray matter surface, average T2 intensity white matter surface, difference between T2 white and gray matter.

A high angular resolution diffusion imaging (HARDI) scan was done to segment white matter tracts and measure diffusion parameters. We used fractional anisotropy (FA) of major white matter tracts labelled by AtlasTrack [35]. FA quantifies the degree of anisotropy, or directionality, of water diffusion, providing insights into the structural integrity and organization of white matter tracts [36]. The processing steps are explained in Hagler and colleagues [8].

Supplementary Figure 1 details all 81 sets of features and their combination in the stacking layer.

#### Cognitive Prediction Models

Within each fold, we standardized the training set features across participants, then applied training statistics (including mean and standard deviation) to normalize the test set before training the predictive model. Within each training set, we performed an additional five-fold cross-validation to optimize the hyperparameters of the predictive models with the negative mean squared error (NMSE) as the evaluation metric.

In our approach, we first built unimodal models by using Partial Least Squares (PLS) to predict cognitive-functioning scores from each of the 81 sets of features. Once these models were trained on the training set fold (with cross-validation for tuning), we computed predicted values from each model. We combined them into 11 different stacking configurations (e.g., stacked all, task contrasts, etc.) and trained a Random Forest model to predict cognitive abilities from each stacked model. For example, to build a stacked model combining task contrasts for ABCD, we first developed unimodal models for each of the contrast-related sets of features (each including 379 or 148 features), and then used the predicted values from these models as individual features in the stacked task contrasts model. This approach, often called late fusion, contrasts with methods that simply concatenate raw features (referred to as early fusion or flat model), and it enables the use of different machine learning algorithms across the various Phenotypes [37].

Partial Least Squares Regression (PLS) is a dimensionality reduction technique that identifies latent components to maximize the covariance between predictor variables X and the response variable Y, solving for components t and u such that $\max \mathrm{Cov}(t,u)=wTX$ and $t = Xw$, $u=Yq$, where w and q are weights for X and Y, respectively [38]. To optimize the number of components, we performed a grid search with five-fold cross-validation, considering all components for sets with fewer than 30 features and capping the number at 30 for larger sets of features. The optimal model was selected based on the negative mean squared error (NMSE) and was used to transform the feature space and generate predictions, reducing dimensionality while retaining predictive information [38].

Random Forest regression is an ensemble learning method that aggregates predictions from multiple decision trees to improve predictive performance and reduce overfitting. Its ability to handle non-linear relationships and robustly deal with missing data made it suitable for the stacking layer. Each tree in the forest is trained on a bootstrapped sample of the data, and at each split, a random subset of features is considered, introducing randomness and diversity [39]. The hyperparameters used include the number of decision trees in the forest, set to 1000, the maximum depth of each tree, searched over a range of 1 to 10 to control model complexity, and the number of features considered at each split, with options including the total number of features (None), the square root of the feature count (sqrt), and the logarithm base 2 of the feature count (log2).

**Evaluation.** Pearson's Correlation ($r$), coefficient of determination ($R^2$), mean absolute error (MAE) were reported as measures of predictive performance across both folds for each training strategy.

| Stacked | Single | N features | N participants | Source |
|---|---|---|---|---|
| | Glasser Nback 2-0back | 379 | 5709 | ABCC |
| | Glasser Nback 2back | 379 | 5709 | ABCC |
| | Glasser Nback 0back | 379 | 5709 | ABCC |
| | Glasser Nback face | 379 | 5709 | ABCC |
| | Glasser Nback emotionface | 379 | 5709 | ABCC |
| | Glasser Nback place | 379 | 5709 | ABCC |
| | Glasser Nback face-place | 379 | 5709 | ABCC |
| | Glasser Nback emotionface-neutralface | 379 | 5709 | ABCC |
| | Glasser Nback NegFace-NeutFace | 379 | 5709 | ABCC |
| | Glasser Nback PosFace-NeutFace | 379 | 5709 | ABCC |
| | Destriuex Nback emotionface-neutralface | 167 | 7716 | ABCD |
| | Destriuex Nback 2back | 167 | 7716 | ABCD |
| | Destriuex Nback face-place | 167 | 7716 | ABCD |
| | Destriuex Nback NegFace-NeutFace | 167 | 7716 | ABCD |
| | Destriuex Nback emotionface | 167 | 7716 | ABCD |
| | Destriuex Nback PosFace-NeutFace | 167 | 7716 | ABCD |
| | Destriuex Nback 0back | 167 | 7716 | ABCD |
| | Destriuex Nback place | 167 | 7716 | ABCD |
| | Destriuex Nback 2-0back | 167 | 7716 | ABCD |
| | Glasser MID Loss-Neutral | 379 | 6777 | ABCC |
| | Glasser MID Reward-Neutral | 379 | 6777 | ABCC |
| | Glasser MID LargeReward-Neutral | 379 | 6777 | ABCC |
| | Glasser MID LargeLoss-Neutral | 379 | 6777 | ABCC |
| | Glasser MID SmallLoss-Neutral | 379 | 6777 | ABCC |
| | Glasser MID LossHit-LossMiss | 379 | 6777 | ABCC |
| | Glasser MID SmallReward-Neutral | 379 | 6777 | ABCC |
| | Glasser MID LargeReward-SmallReward | 379 | 6777 | ABCC |
| | Glasser MID LargeLoss-SmallLoss | 379 | 6777 | ABCC |
| | Glasser MID RewardHit-RewardMiss | 379 | 6777 | ABCC |
| | Destriuex MID LargeReward-SmallReward | 167 | 9102 | ABCD |
| | Destriuex MID LossHit-LossMiss | 167 | 9102 | ABCD |
| | Destriuex MID Reward-Neutral | 167 | 9102 | ABCD |
| | Destriuex MID SmallLoss-Neutral | 167 | 9102 | ABCD |
| | Destriuex MID LargeLoss-Neutral | 167 | 9102 | ABCD |
| | Destriuex MID SmallReward-Neutral | 167 | 9102 | ABCD |
| | Destriuex MID LargeReward-Neutral | 167 | 9102 | ABCD |
| | Destriuex MID Loss-Neutral | 167 | 9102 | ABCD |
| | Destriuex MID LargeLoss-SmallLoss | 167 | 9102 | ABCD |
| | Destriuex MID RewardHit-RewardMiss | 167 | 9102 | ABCD |
| | Glasser SST Stop-CorrectGo | 379 | 5998 | ABCC |
| | Glasser SST IncorrectStop-CorrectGo | 379 | 5998 | ABCC |
| | Glasser SST IncorrectGo | 379 | 5998 | ABCC |
| | Glasser SST IncorrectStop | 379 | 5998 | ABCC |
| | Glasser SST IncorrectGo-CorrectGo | 379 | 5998 | ABCC |
| | Glasser SST CorrectStop | 379 | 5998 | ABCC |
| | Glasser SST CorrectStop-CorrectGo | 379 | 5998 | ABCC |
| | Glasser SST IncorrectGo-IncorrectStop | 379 | 5998 | ABCC |
| | Glasser SST CorrectGo | 379 | 5998 | ABCC |
| | Glasser SST CorrectStop-IncorrectStop | 379 | 5998 | ABCC |
| | Destriuex SST IncorrectGo-IncorrectStop | 167 | 8034 | ABCD |
| | Destriuex SST AnyStop-CorrectStop | 167 | 8034 | ABCD |
| | Destriuex SST IncorrectStop-CorrectGo | 167 | 8034 | ABCD |
| | Destriuex SST IncorrectGo-CorrectGo | 167 | 8034 | ABCD |
| | Destriuex SST CorrectGo-fixation | 167 | 8034 | ABCD |
| | Destriuex SST CorrectStop-IncorrectStop | 167 | 8034 | ABCD |
| | Destriuex SST CorrectStop-CorrectGo | 167 | 8034 | ABCD |

| | | | |
|---|---|---|---|
| Glasser Nback FC | 71631 | 4511 | ABCC |
| Glasser MID FC | 71631 | 5067 | ABCC |
| Glasser SST FC | 71631 | 4987 | ABCC |
| Glasser rest FC | 71631 | 5253 | ABCC |
| General FC | 71631 | 4273 | ABCC |
| Multitask FC | 71631 | 5027 | ABCC |
| rsfMRI subcortical-network FC | 247 | 9299 | ABCD |
| rsfMRI cortical FC | 91 | 9299 | ABCD |
| rsfMRI temporal variance | 352 | 9299 | ABCD |
| T2 gray matter avg intensity | 148 | 10471 | ABCD |
| T2 white matter avg intensity | 148 | 10471 | ABCD |
| T2 normalised intensity | 148 | 10471 | ABCD |
| T1 normalised intensity | 148 | 11177 | ABCD |
| T1 gray matter avg intensity | 148 | 11177 | ABCD |
| T1 white matter avg intensity | 148 | 11177 | ABCD |
| T2 summations | 4 | 10471 | ABCD |
| T1 summations | 9 | 11177 | ABCD |
| T1 Subcortical Volume | 19 | 11177 | ABCD |
| T2 Subcortical Volume | 19 | 10471 | ABCD |
| Total brain volume | 5 | 9141 | ABCC |
| Surface area | 148 | 9141 | ABCC |
| Sulcal depth | 148 | 11177 | ABCD |
| Cortical thickness | 148 | 9141 | ABCC |
| Cortical volume | 148 | 11177 | ABCD |
| DTI | 23 | 10194 | ABCD |

| |
|---|
| Stacked all |
| Task FCs + Contrasts |
| Task Contrasts |
| Non-Task |
| Task FCs |
| Nback Contrasts |
| MID Contrasts |
| SST Contrasts |
| Rest FCs |
| All FCs |
| sMRI |

**Supplementary Figure 1 Unimodal and multimodal phenotypes**.

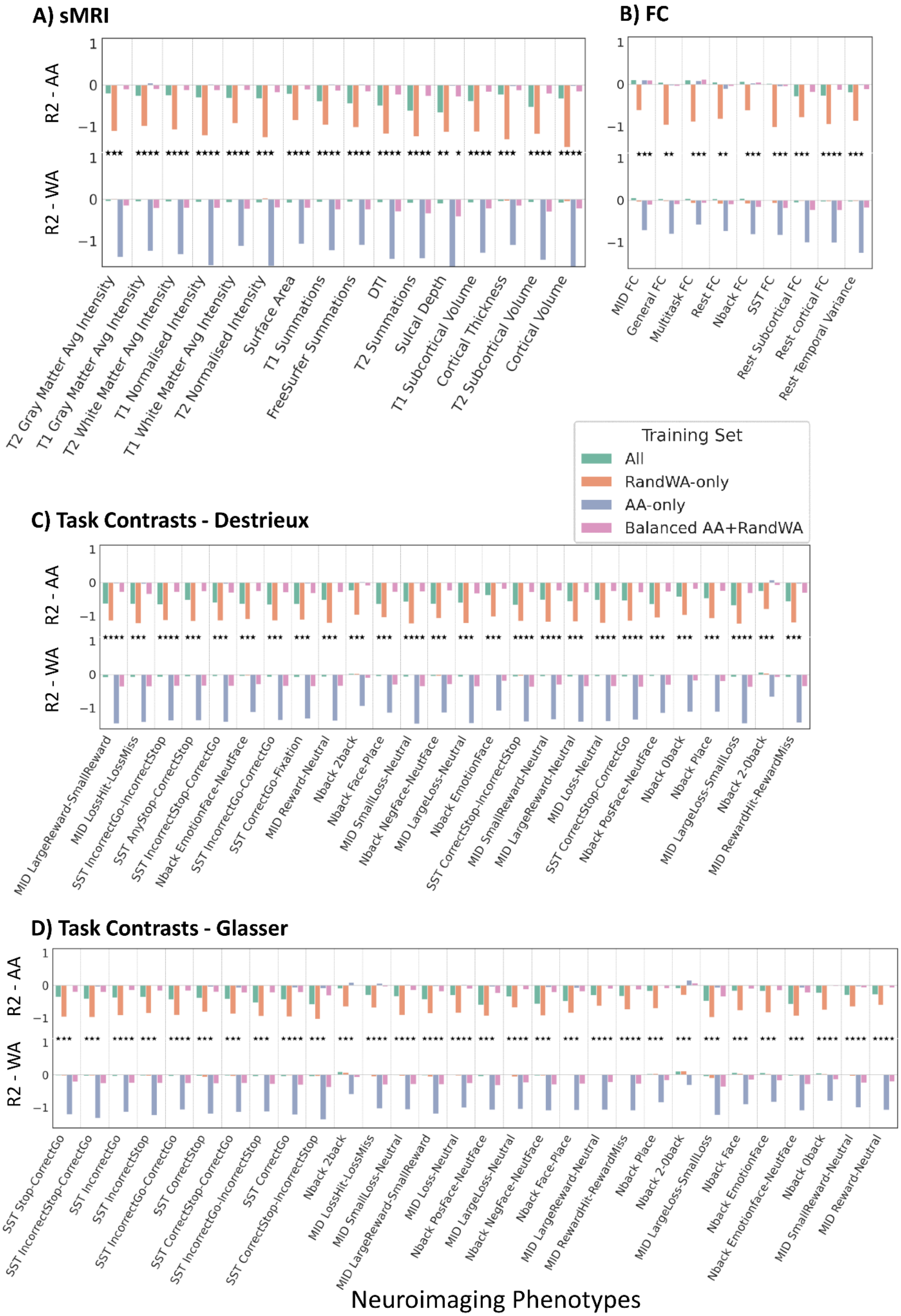
A) sMRI
B) FC
C) Task Contrasts - Destrieux
D) Task Contrasts - Glasser
R2 - AA
R2 - WA
T2 Gray Matter Avg Intensity
T1 Gray Matter Avg Intensity
T2 White Matter Avg Intensity
T1 Normalised Intensity
T1 White Matter Avg Intensity
T2 Normalised Intensity
Surface Area
T1 Summations
FreeSurfer Summations
DTI
T2 Summations
Sulcal Depth
T1 Subcortical Volume
Cortical Thickness
T2 Subcortical Volume
Cortical Volume
MID FC
General FC
Multitask FC
Rest FC
Nback FC
SST FC
Rest Subcortical FC
Rest cortical FC
Rest Temporal Variance
Training Set
All
RandWA-only
AA-only
Balanced AA+RandWA
MID LargeReward-SmallReward
MID LossHit-LossMiss
SST IncorrectGo-IncorrectStop
SST AnyStop-CorrectStop
SST IncorrectStop-CorrectGo
Nback EmotionFace-NeutFace
SST IncorrectGo-CorrectGo
SST CorrectGo-Fixation
MID Reward-Neutral
Nback 2back
Nback Face-Place
MID SmallLoss-Neutral
Nback NegFace-NeutFace
MID LargeLoss-Neutral
Nback EmotionFace
SST CorrectStop-IncorrectStop
MID SmallReward-Neutral
MID LargeReward-Neutral
MID Loss-Neutral
SST CorrectStop-CorrectGo
Nback PosFace-NeutFace
Nback 0back
Nback Place
MID LargeLoss-SmallLoss
Nback 2-0back
MID RewardHit-RewardMiss
SST Stop-CorrectGo
SST IncorrectStop-CorrectGo
SST IncorrectGo
SST IncorrectStop
SST IncorrectGo-CorrectGo
SST CorrectStop
SST CorrectStop-CorrectGo
SST IncorrectGo-IncorrectStop
SST CorrectGo
SST CorrectStop-IncorrectStop
Nback 2back
MID LossHit-LossMiss
MID SmallLoss-Neutral
MID LargeReward-SmallReward
MID Loss-Neutral
Nback PosFace-NeutFace
MID LargeLoss-Neutral
Nback NegFace-NeutFace
Nback Face-Place
MID LargeReward-Neutral
MID RewardHit-RewardMiss
Nback Place
Nback 2-0back
MID LargeLoss-SmallLoss
Nback Face
Nback EmotionFace
Nback Emotionface-NeutFace
Nback 0back
MID SmallReward-Neutral
MID Reward-Neutral
Neuroimaging Phenotypes

**Supplementary Figure 2 Coefficient of Determination (R2) for Test AA and WA Participants Across Different Phenotypes: (A) sMRI and DTI, (B) FC, (C) Task Contrast Parcellated with Glasser, and (D) Task Contrast Parcellated with Destrieux.** Stars indicate significantly different performance between tests AA and WA based on the permutation tests (*p-value* = 0.05)**.** *Sulcal Depth was excluded from main analysis due to unstable model performance.

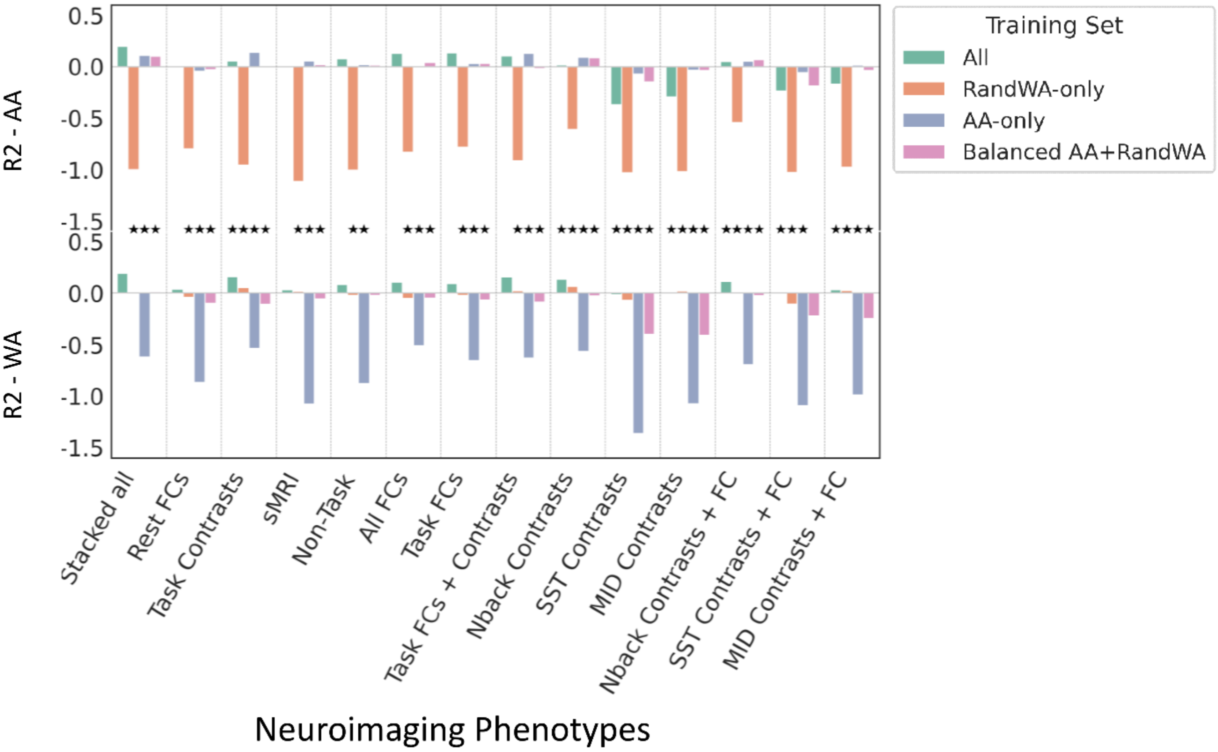


**Supplementary Figure 3 Coefficient of Determination (R2) for Test AA and WA Participants Across Stacked Multimodal Phenotypes.** Stars indicate significantly different performance between tests AA and WA based on the permutation tests (*p-value* = 0.05)**.**

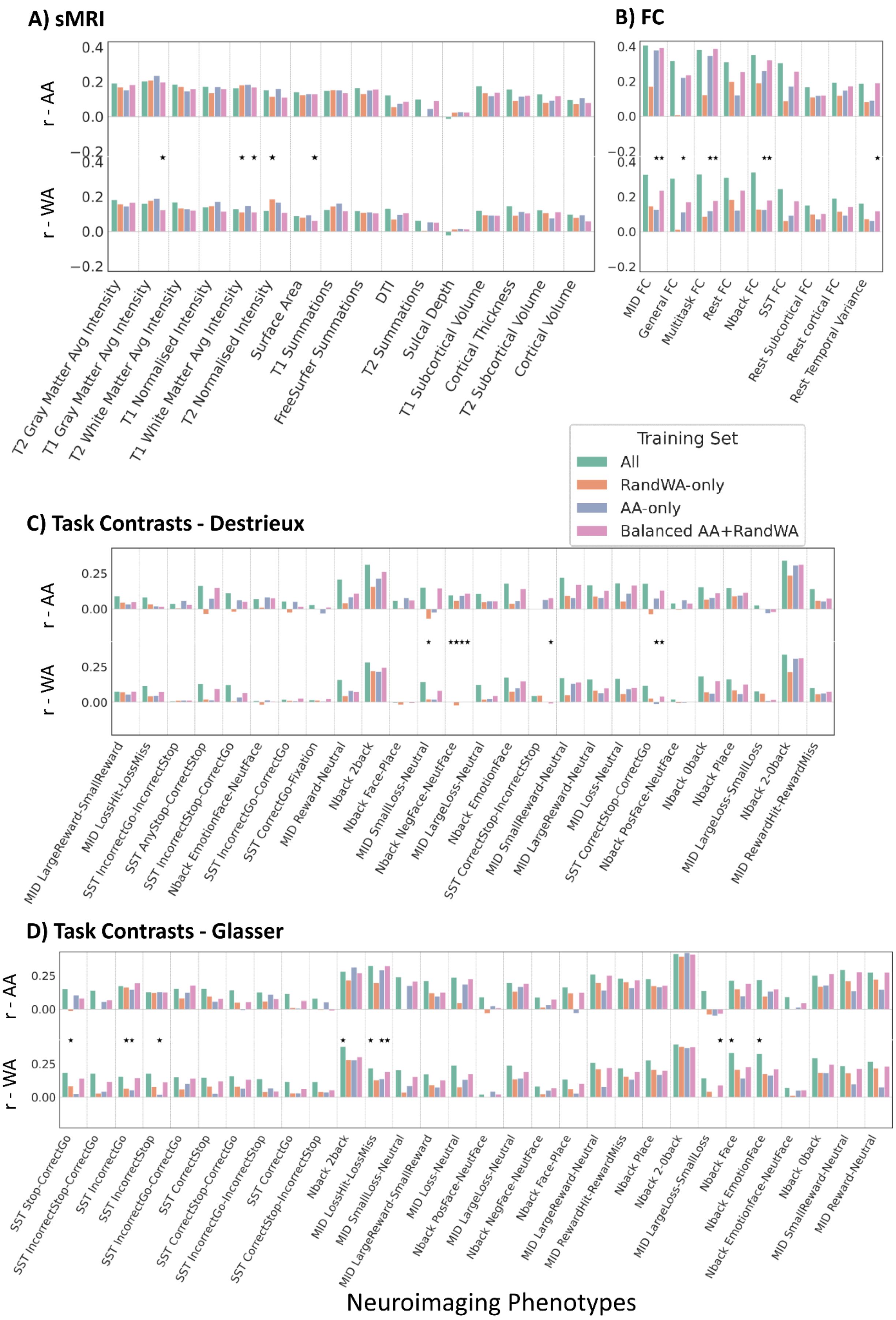
A) sMRI
B) FC
r - AA
r - WA
0.4
0.2
0.0
−0.2
T2 Gray Matter Avg Intensity
T1 Gray Matter Avg Intensity
T2 White Matter Avg Intensity
T1 Normalised Intensity
T1 White Matter Avg Intensity
T2 Normalised Intensity
Surface Area
T1 Summations
FreeSurfer Summations
DTI
T2 Summations
Sulcal Depth
T1 Subcortical Volume
Cortical Thickness
T2 Subcortical Volume
Cortical Volume
MID FC
General FC
Multitask FC
Rest FC
Nback FC
SST FC
Rest Subcortical FC
Rest cortical FC
Rest Temporal Variance
Training Set
All
RandWA-only
AA-only
Balanced AA+RandWA
C) Task Contrasts - Destrieux
0.25
0.00
MID LargeReward-SmallReward
MID LossHit-LossMiss
SST IncorrectGo-IncorrectStop
SST AnyStop-CorrectStop
SST IncorrectStop-CorrectGo
Nback EmotionFace-NeutFace
SST IncorrectGo-CorrectGo
SST CorrectGo-Fixation
MID Reward-Neutral
Nback 2back
Nback Face-Place
MID SmallLoss-Neutral
Nback NegFace-NeutFace
MID LargeLoss-Neutral
Nback EmotionFace
SST CorrectStop-IncorrectStop
MID SmallReward-Neutral
MID LargeReward-Neutral
MID Loss-Neutral
SST CorrectStop-CorrectGo
Nback PosFace-NeutFace
Nback 0back
Nback Place
MID LargeLoss-SmallLoss
Nback 2-0back
MID RewardHit-RewardMiss
D) Task Contrasts - Glasser
SST Stop-CorrectGo
SST IncorrectStop-CorrectGo
SST IncorrectGo
SST IncorrectStop
SST IncorrectGo-CorrectGo
SST CorrectStop
SST CorrectStop-CorrectGo
SST IncorrectGo-IncorrectStop
SST CorrectGo
SST CorrectStop-IncorrectStop
Nback 2back
MID LossHit-LossMiss
MID SmallLoss-Neutral
MID LargeReward-SmallReward
MID Loss-Neutral
Nback PosFace-NeutFace
MID LargeLoss-Neutral
Nback NegFace-NeutFace
Nback Face-Place
MID LargeReward-Neutral
MID RewardHit-RewardMiss
Nback Place
Nback 2-0back
MID LargeLoss-SmallLoss
Nback Face
Nback EmotionFace
Nback Emotionface-NeutFace
Nback 0back
MID SmallReward-Neutral
MID Reward-Neutral
Neuroimaging Phenotypes

**Supplementary Figure 4 Pearson's correlation (r) for Test AA and WA Participants Across Different Phenotypes: (A) sMRI and DTI, (B) FC, (C) Task Contrast Parcellated with Glasser, and (D) Task Contrast Parcellated with Destrieux.** Stars indicate significantly different performance between tests AA and WA based on the permutation tests (*p-value* = 0.05)**.**

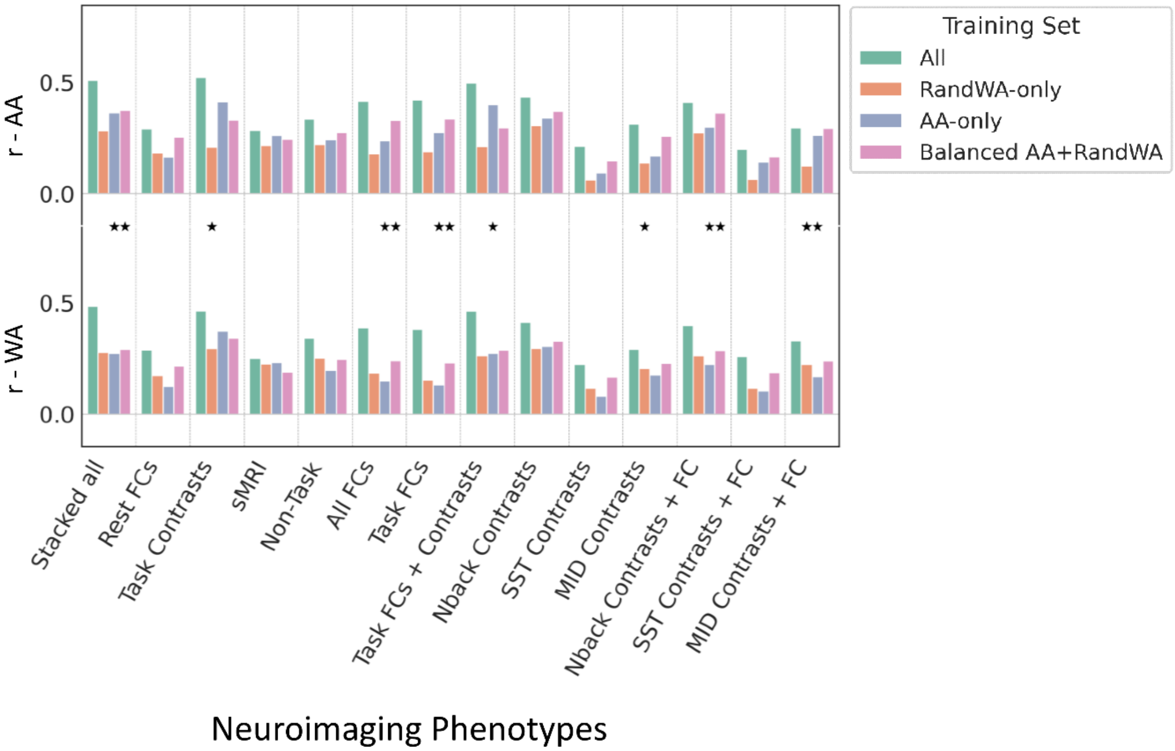


**Supplementary Figure 5 Pearson's correlation (r) for Test AA and WA Participants Across Different Stacked Multimodal Phenotypes.** Stars indicate significantly different performance between tests AA and WA based on the permutation tests (*p-value* = 0.05)**.**
